# Genomic changes are varied across congeneric species pairs of animals

**DOI:** 10.1101/2024.09.05.611358

**Authors:** Warren R. Francis, Sergio Vargas, Gert Wörheide

## Abstract

Synteny, the shared arrangement of genes on chromosomes between related species, is a marker of shared ancestry, and synteny-breaking events can result in genomic incompatibilities between populations and ultimately lead to speciation events. Despite its pivotal role as a driver of speciation, the role of synteny breaks on speciation is poorly studied due to a lack of chromosome-level genome assemblies for a taxonomically broad sample of organisms. Here, using 22 congeneric animal genome pairs, we find a link between protein identity, microsynteny, and macrosynteny, but no evidence for a universal path of genomic change during divergence. We observed varied trajectories of synteny conservation relative to protein identity in non-model organisms, with many species pairs showing no karyotypic changes and others displaying large genomic rearrangements. This contrasts with previous studies on model organisms and indicates that the genomic changes preceding or resulting from speciation are likely very contextual between clades.

## Introduction

The “species” is a fundamental concept in biology, and many different concepts – at least 26 – exist of what constitutes a species and how species should be defined and delineated (De Queiroz, 2007; Hausdorf, 2011; Zachos, 2016). Delimiting species and correctly assessing species numbers in an ecosystem is essential to assess, for example, the resilience of species assemblages and ecosystems to environmental challenges, with consequences for biodiversity conservation and management (Frankham et al., 2012; Hey et al., 2003). Probably the most important species concept proposed is the “biological species” of Ernst Mayr (Mayr, 1942). Mayr defined species as “*groups of actually or potentially interbreeding natural populations, which are reproductively isolated from other such groups*”. This condition is only testable for (a few) species that can be bred under controlled conditions, for example, domesticated species or organisms that can be easily kept in the laboratory (rodents, flowers, etc.). However, it may be very difficult or impossible to directly test for most organisms, especially marine species. Hence, other “more practical” species concepts, such as the phylogenetic species concept, the evolutionary species concept, and the morphological species concept, are frequently used (Frankham et al., 2012; reviewed in Ghiselin, 2010).

Under the biological species concept, reproductive isolation is key. This is because even if the exact mechanisms leading to the reproductive isolation of two or more populations remain unknown, it is predicted that reproductively isolated populations will become genetically incompatible with time through the accumulation of random mutations (i.e., Dobzhansky-Müller incompatibilities) in their genomes (Orr, 1996). Importantly, this process can occur if the populations are under selection or can proceed under pure genetic drift. Thus, the accumulation of Dobzhansky-Müller incompatibilities provides a universal mechanism whereby species could arise in nature. For instance, genetic incompatibilities leading to hybrid inviability have been detected in mice (Vrana et al., 2000; Zechner et al., 1996), and in shrimp sperm/eggs recognition incompatibility has been described (Ulate & Alfaro-Montoya, 2010). However, the genomic causes leading to genetic incompatibility and, eventually, hybrid inviability remain to be assessed for most species.

Although the Dobzahnsky-Müller model typically considers mutations as the causal agent behind genetic incompatibilities, synteny-breaking genome rearrangements could have similar effects on hybrid viability. In this regard, changes in both macro-synteny (i.e., the shared gene content on a chromosome, Figure 1A) after chromosomal translocations, for instance, and micro-synteny (i.e., the preserved order of several genes in a chromosome, Figure 1B) can lead to chromosome pairing problems during mitosis in hybrid zygotes, resulting in hybrid inviability and, eventually, in speciation. Several studies have observed that macro-synteny is well-conserved across animal phyla (Kenny et al., 2020; Lv et al., 2011; Simakov et al., 2022; Srivastava et al., 2008) suggesting that chromosome gene content is constrained and that translocations are either infrequent or selected against across long time scales. One proposed mechanism underlying this constraint is that gene dosage of multiple genes on a chromosome must be maintained (Lv et al., 2011). Since chromosome translocations may result in a zygote/cell with either 1x or 3x copies of a gene, it is likely that the loss of viability resulting from these chromosome rearrangements is selected negatively, as proposed by Ohno (1970). Thus, translocations may only play a minor role in speciation as their effects are too deleterious to spread in a population while other macro-synteny changes, such as Robertsonian fusions, may still have effects on speciation.

**Figure 1:**
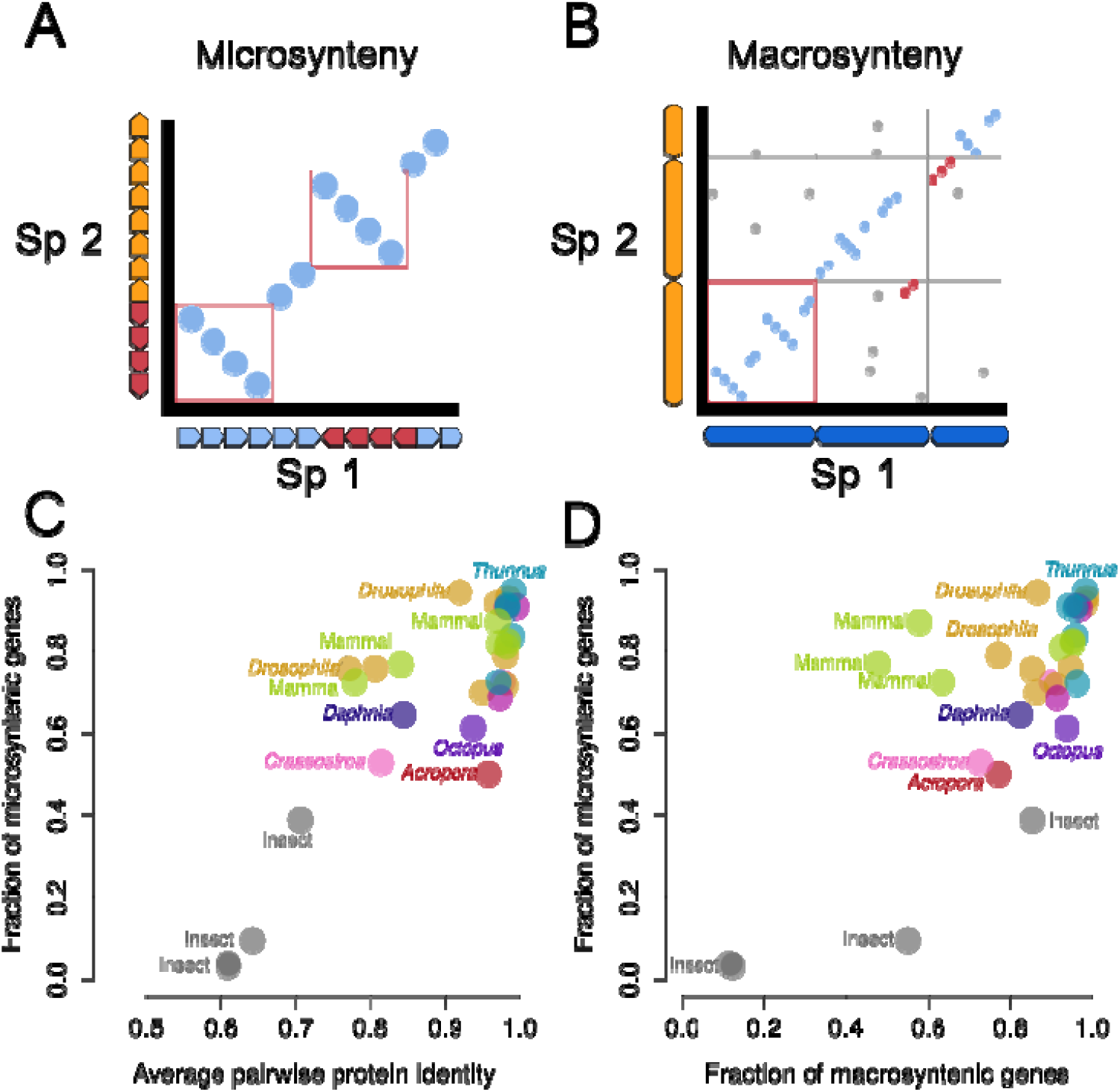
(A) Schematic dot plot of microsynteny. Arrows along the axes represent genes, with red genes indicating inversions in one of the two genomes. Red boxes indicate microsyntenic blocks. (B) Schematic dot plot of macrosynteny. Bars along the axes represent chromosomes and points represent positions of homologous genes between the two species. Blue points would be considered macrosyntenic, red points are microsyntenic but with a translocation, and gray points are any other hits, e.g. genes sitting on very different positions in the compared species pairs. (C) Pairwise protein sequence identity for all orthologs plotted against fraction of genes in synteny blocks. Gray (insects) and yellow-green (mammals) indicate additional comparisons between species not in the same genus. The dotted line indicates the calculated regression line from (Zdobnov & Bork, 2007) of the same two axes. (D) Relationship between microsynteny and macrosynteny.

In contrast, changes in micro-synteny resulting from chromosome fusions and inversions appear to be more widespread, possibly because they do not result in gene dosage imbalances. This is expected to some extent because eukaryotic genes typically have their own promoters so that genes “moving” together with their linked regulatory regions should preserve their expression patterns. Supporting this idea, the expression of nested genes appears to be uncorrelated, and gene proximity and dosage typically are also uncorrelated (Assis et al., 2008). There are some exceptions: genes controlled by bidirectional promotors or highly-conserved non-coding elements (HCNEs) controlling a group of neighbouring genes (Engström et al., 2007; Kikuta et al., 2007); or genes with regulatory elements inside the introns of neighbouring genes (e.g., the human lactase, which enhancer is located inside its upstream gene *mcmc6* (Enattah et al., 2002)). Thus, it is likely that micro-synteny changes are either neutral or only slightly deleterious for most eukaryotic genes allowing these changes to spread in populations after their appearance and eventually become fixed (Ohno, 1970; Slijepcevic, 1998).

Despite the potential role of micro-synteny in speciation, the degree of micro-synteny is variable within the animal lineages examined to date. For instance, extensive syntenic blocks, of tens to hundreds of genes, have been found in octocorals (Jiang et al., 2019) but not medusozoans (Jiang et al., 2019; Khalturin et al., 2019), or in glass sponges (Francis et al., 2023) but not demosponges (Kenny et al., 2020). Even in two smaller phyla, Placozoa and Ctenophora, microsynteny was extensive in placozoans (Eitel et al., 2018), but was almost completely absent in ctenophores (Schultz et al., 2021). The effects of conserved synteny have only been examined in model organisms and their close relatives, making it difficult to generalise the role of micro-synteny breaking in speciation across the animal kingdom. To evaluate this, we examined macro- and micro-synteny in 22 con-generic pairs with chromosome-level genome assemblies representing five animal phyla. We find varied trajectories of synteny conservation, suggesting that no universal path of genomic change during speciation exists in animals.

## Material & Methods

### Datasets from NCBI

A total of 17 animal genera accounting for 22 pairs of congeneric species were downloaded from NCBI (see Table 1, and Supplemental Figure 1). The bulk of the 55 available congeneric genome pairs on NCBI mostly consisted of arthropods (e.g., 18 *Drosophila* species) and chordates (mostly mammals, e.g., cow, pig, mouse). We limited this selection to include only chromosome-level genome assemblies for which NCBI annotations for that specific version were also available, and also excluded redundant pairs within genera from the analyses e.g. we only kept 4 *Drosophila* species (see Table 1). In one of the genera, *Crassostrea*, two of the species were reassigned into another genus *Magallana* (Salvi & Mariottini, 2017) on NCBI during the review of this manuscript, though the original comparisons are retained below. For additional comparisons to remake the analyses of previous studies, we analyzed the insects against each other, using *Drosophila melanogaster* as one species, and then one of *Anastrepha, Culex, Bombus*, and *Vespa* for each pair (matching what was done in (Zdobnov & Bork, 2007). We also included a set of seven genomes of additional mammal genera, namely human, *Mus musculus* (mouse), *Pan troglodytes* (chimpanzee), *Gorilla gorilla* (gorilla), *Pongo abelii* (orangutan), *Nomascus leucogenys* (gibbon), and *Monodelphis domestica* (opossum).

**Table 1:** The 17 genera used in analysis. ^a^ several unassigned scaffolds in both *Bombus* species appear roughly the size of chromosomes, and would account for an extra 4-6 chromosomes. ^b^ at the time of downloading the data from NCBI, all species used were of the genus *Crassostrea*, though now some species in this genus have been reassigned to *Magellana* (see Salvi & Mariottini, 2017, 2021) ^c^ 6Mb of unassigned scaffolds are given as a single scaffold in *V. crabro*, with mostly masked bases and effectively no annotated genes, hence no synteny was detected with *V. velutina*.

| Genus | N spp. | Representing taxonomic group | Phylum | Genome size (Mb) | N Chrs |
| --- | --- | --- | --- | --- | --- |
| *Acropora* | 2 | Staghorn corals | Cnidaria | 446-475 | 14 |
| *Anastrepha* | 2 | Fruit flies | Arthropoda | 820-836 | 6 |
| *Bombus* | 2 | Bumblebees | Arthropoda | 254-392 | 18 (24 <sup>a</sup>) |
| *Bufo* | 2 | Toads | Chordata | 4545-5044 | 11 |
| *Cervus* | 2 | Deer | Chordata | 2526-2886 | 34 |
| *Crassostrea/*<br>*Magallana* <sup>(b)</sup> | 3 | Oysters | Mollusca | 624-684 | 10 |
| *Culex* | 2 | Mosquitoes | Arthropoda | 566-573 | 3 |
| *Daphnia* | 2 | Water fleas | Arthropoda | 133-161 | 10,12 |
| *Drosophila* | 4 | Fruit flies | Arthropoda | 143-191 | 4-7 |
| *Epinephelus* | 3 | Groupers (fish) | Chordata | 1030-1087 | 24 |
| *Lytechinus* | 2 | Sea urchins | Echinodermata | 869-999 | 19 |
| *Mauremys* | 2 | Turtles | Chordata | 2367-2484 | 27 |
| *Octopus* | 2 | Octopus | Mollusca | 2342-2719 | 30 |
| *Oreochromis* | 2 | Tilapia (fish) | Chordata | 1005 | 22 |
| *Perca* | 2 | Perch (fish) | Chordata | 877-951 | 24 |
| *Thunnus* | 2 | Tuna (fish) | Chordata | 782-792 | 24 |
| *Vespa* | 2 | Hornets | Arthropoda | 193-229 | 25 (26 <sup>c</sup>) |

### Addition of representatives from a coral genus

For the coral genus *Acropora*, both *A. hyacinthus* and *A. millepora* had chromosome-level assemblies, but only *A. millepora* had an annotation file on NCBI. Thus, we used the annotation file from the previous version of *A. hyacinthus*, downloaded from the Marine Genomics Group at OIST (https://marinegenomics.oist.jp/ahya/viewer/download?project_id=91). From the v1 annotation (Shinzato et al., 2021), we generated transcripts using gffread v0.12.7 (Pertea & Pertea, 2020), which were then filtered to remove splice variants, and mapped to the chromosome-level assembly using minimap2 v2.23 (Li, 2018).

### Removal of additional protein isoforms

Because the NCBI annotation pipeline will include splice variants where evidence is available, several pre-processing steps were applied for each species to avoid isoforms from interfering with nearly all steps of the analysis (e.g., counting duplicate orthologs). For our analyses, only the longest isoform was retained for each gene using a custom Python script “get_genbank_longest_isoforms.py”. This script simultaneously removes the shorter protein isoforms for each gene and the annotation entries from the respective GFF files. It should be noted that the longest isoform is not necessarily the “primary” or “correct” one, however, the role of splice variation in evolution or phylogenetics is not well-studied, and the main focus of our analyses was the maintenance or break of synteny blocks that can be identified with any of the alternative isoforms. The filtered versions of the genomes were used for all subsequent analyses, including the microsynteny and protein sequence identity. Scripts are available online at https://github.com/PalMuc/congeneric_synteny/tree/main.

### Generation of ortholog pairs

For each species pair, homologous protein clusters were generated using the following strategy. Proteins were aligned using DIAMOND v2.0.13 (Buchfink et al., 2015) with default parameters. Ortholog pairs were generated using the custom script ‘makehomologs.py’, which first filters the DIAMOND hits by removing self-matches, removing matches with a low bitscore to length ratio, and finally leverages ‘MCL’ for the actual clustering (Enright et al., 2002).

Several filtering steps were applied (see Supplemental Figure 1), and only one-to-one protein clusters were retained for subsequent analysis (i.e., paralogs were removed). This means that fragmented proteins (where one gene is fragmented in either genome and results in two proteins that align to a single protein in the other genome) were excluded. This also means that genes duplicated in either genome (1-to-many and many-to-many orthologs) are also excluded, even if they were identical. It should be noted that this method does not directly identify pairs by synteny. That is, clustered proteins or identical proteins may still be omitted if that cluster is not single-copy in both genomes, even if the synteny potentially could be detected by a stricter approach. For instance, the three copies of human calmodulin are 100% identical to each other and to the four mouse homologs, but this would result in a cluster of seven proteins, and then get excluded from the analysis regardless of their chromosomal arrangement. Each ortholog pair was then subsequently aligned with MAFFT v7.487 (Katoh & Standley, 2013) to account for the entire protein in calculations of protein identity. Amino acid sites were defined as identical, different, or gaps for each species using the custom Python script “alignment_conserved_site_to_dots.py”. Protein identity was calculated as the number of identities divided by the number of identities plus differences, meaning that gap sites were ignored. The frequency of gaps by species is examined in Supplemental Figure 2, but does not appear to cause any obvious bias, such as if highly-divergent regions were systematically missed in many protein pairs.

### Quantification of microsynteny in genome pairs

We then quantified the number of genes that could be found in syntenic blocks between each pair of congeneric species using the custom Python script “microsynteny.py”, with various options depending on the species pair (see https://github.com/wrf/genomeGTFtools/blob/master/microsynteny.py). Here, we define microsynteny as blocks of three or more consecutive genes in the same order in both species, which are then assumed to be homologous, or “identical by descent”.

As intergenic length is linearly related to total genome size (Francis & Wörheide, 2017), programs measuring microsynteny are very sensitive to any parameters controlling the allowed distance to the next gene. We calibrated the allowed distance to the next gene across several species, finding the optimal length of ca. 1/5000 of the total length of the genome (Supplemental Figure 3). For instance, for the tuna, this length accounted for 99.4% of intergenic distances (Supplemental Figure 4), meaning that a value was used to account for nearly all genes. Identifying syntenic blocks can also be affected by gene insertions or deletions, so the minimum block length and number of “intervening genes” (i.e., those inserted/deleted) were also evaluated. Based on this, we required a minimum of three genes per block and allowed up to four intervening genes (Supplemental Figure 5). When the minimum was set to two genes per block, as was used by (Zdobnov & Bork, 2007), the results were not appreciably different. Since spurious synteny blocks were frequent in randomised datasets when the minimum number of genes per block was set to two (Supplemental Figure 5 A, B) and were greatly reduced (10-20 fold reduction) if the minimum number of syntenic genes was set to three (Supplemental Figure 5 A) we used three genes per block as a conservative approach to synteny determination.

### Quantification of macrosynteny

For macrosynteny, we counted the number of genes that could be found on homologous chromosomes between each pair of congeneric species using custom scripts. All proteins were aligned using DIAMOND v2.0.13 (Buchfink et al., 2015), and the resulting alignments were used to generate dot plots with the Python script “scaffold_synteny.py”. Macrosynteny was then determined using the R script “calculate_main_block_genes.R” by counting the number of genes for each scaffold that have DIAMOND matches to the most common scaffold in the other species; these genes were counted as “main” or “macrosyntenic” genes. All genes matching any other scaffold were classified as “non-macrosyntenic”. To remove genes from unassigned scaffolds, we only considered scaffolds greater than 1% of the total assembly size. All code is available at: https://github.com/PalMuc/congeneric_synteny/tree/main

## Results and Discussion

### Microsynteny broadly correlates with protein identity across animals

We found a loose correlation between pairwise protein identity and the number of genes in syntenic blocks, with most species pairs differing in synteny but showing relatively high sequence identity (Figure 1C). For instance, the fraction of syntenic genes in the coral genus *Acropora* was 50.2%, but its average protein identity was 95.8%. The reduced synteny at high protein identity levels indicates that in many species, microsynteny is not as selectively constrained as amino acid sequence.

On the other hand, in some genera (notably, *Drosophila melanogaster* against *D. grimshawi*) pairs show low protein identity but are relatively conserved gene order. In this case, sequence identity decreases across the genus without substantial synteny breaks, indicating that synteny may be constrained. Because insertion of new genes (duplications, (retro)transpositions) or deletions would also disrupt synteny, these processes may also be uncommon in *Drosophila*.

To compare to analyses from other studies, we added two sets of species pairs, that included species not in the same genus, one from insects (gray) and one from mammals (yellow-green) (Figure 1C). While there is not any obvious connection between insects and mammals, these additional sets showed a different pattern of protein sequence identity and microsynteny. This suggests a different genomic history, whereby two events occur: one is the overall complete loss of synteny; the other is a decreasing protein identity, such that orthologs can no longer be identified by BLAST (see Supplemental Figure 6). This may be accompanied further by an overall decrease in the total number of ortholog pairs, because our pipeline excludes recently duplicated genes. In general, this agrees with earlier findings in insects (Zdobnov & Bork, 2007), though that study was focused on insect species between orders.

### Microsynteny changes more frequently than macrosynteny

Given that many inversions (of three or more genes) do not disrupt microsynteny, many inverted blocks would still be detected as syntenic, and the slow accumulation of inversions would not change the fraction of genes in blocks or affect the overall microsynteny estimates. Translocations, however, can change macrosynteny but not microsynteny, as genes move as a block to a different chromosome. We examined this for each congeneric species pair as a function of the number of genes on homologous chromosomes in both species compared to the number of translocated genes or inserted novel genes.

Like the relationship with protein identity, we found that macrosynteny and microsynteny broadly tend to correlate together, (Figure 1D), but can be decoupled in many species pairs. In most species pairs (other than the mammal set) microsynteny appears to be less constrained than macrosynteny. For instance, although microsynteny is almost absent in the insects analysed, macrosynteny remains broadly detectable in these organisms. Mammals, however, display a different pattern of chromosomal evolution, with several species’ pairs showing chromosomal rearrangements while preserving their microsynteny.

While there are examples of changes in karyotype between species (such as the fusion resulting in Chr 2 in humans (Yunis & Prakash, 1982)), only a few of the species in our study had obvious karyotype differences. Macrosynteny is relatively conserved among *Drosophila* species with very conserved microsynteny and relatively low protein sequence identity. Between *D. melanogaster* and *D. erecta* (the closest pair in our study), only one translocation is evident in the genome (Figure 2A). In *Octopus*, which possesses the largest invertebrate genome in our study, the karyotype appears to be the same between the two species analysed, but with frequent small inversions in most chromosomes (Figure 2B). Gene arrangement in *Octopus* is known to be variable, such as the rearrangement of the Hox cluster (Albertin et al., 2015). Other species have karyotype variations. The water flea *Daphnia pulex* has 12 chromosomes, while *D. magna* has only ten. These *Daphnia* species show conserved macrosynteny but rapidly decaying microsynteny (Figure 2C). Also, the karyotype change in *Daphnia* appears to have been caused by recent Robertsonian fission/fusion events in one of the species. For *Crassostrea* / *Magallana* (oysters), two opposite chromosomal fusion/fission events were evident (Figure 2D), producing the same number of chromosomes in the two species.

**Figure 2:**
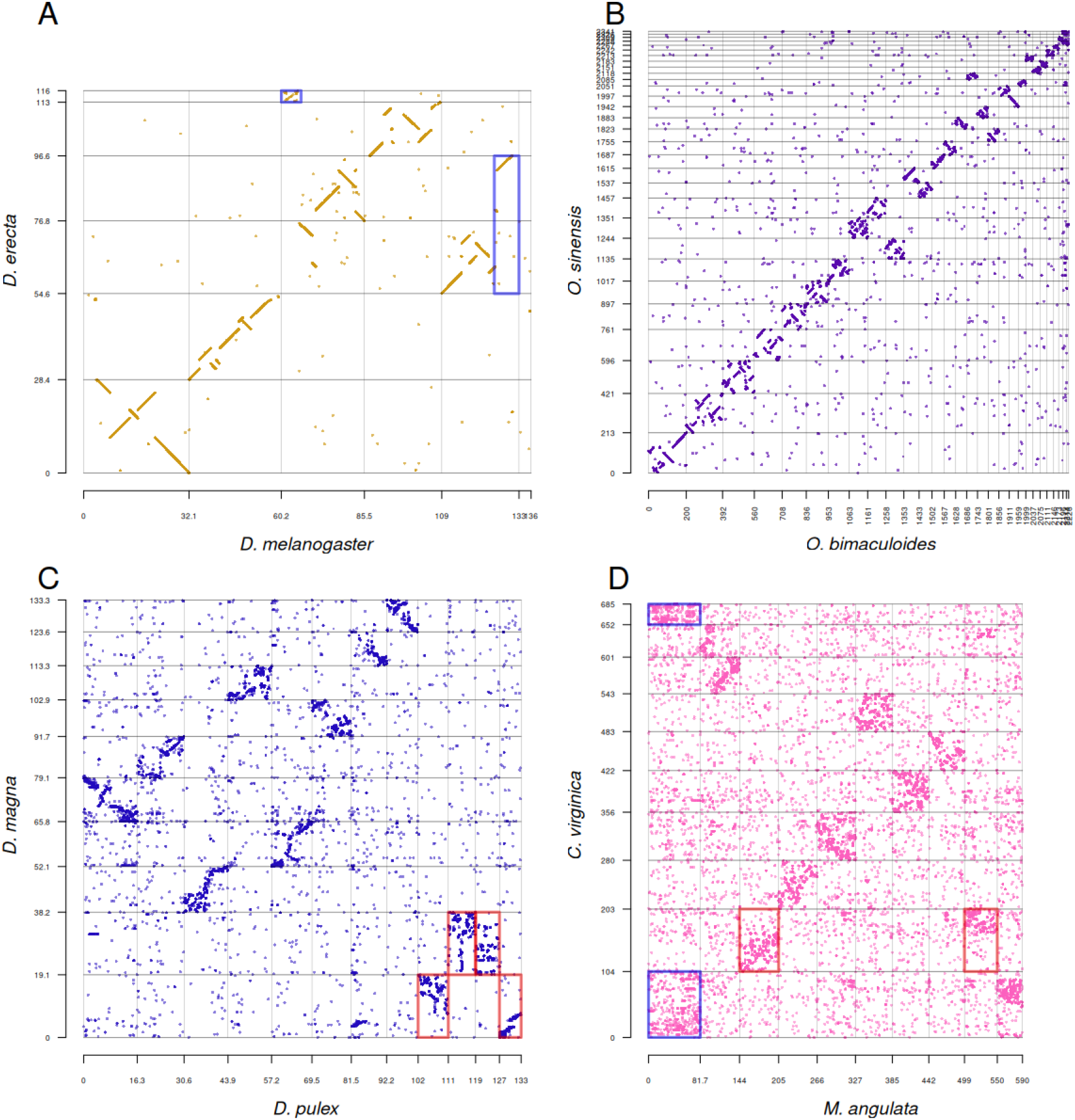
Example dotplots between congeneric species, showing a range of karyotypic states. All axes indicate positions in the genome in megabases. (A) *Drosophila*, showing two translocations/splits in blue boxes, one of which is unassigned to a chromosome in the assembly. (B) *Octopus*, with no obvious translocations but many small inversions. (C) *Daphnia*, red boxes indicate two chromosomal fusions/fissions with mixing in *D. magna*. (D) *Crassostrea/Magallana*, with two evident chromosomal fusions/fissions and split between the two species, as red and blue boxes, respectively. See online repository data for higher resolution versions of all species pairs.

### Distribution of protein identity values is varied across species pairs

We then examined genomic signatures of speciation across the pairs of congeneric species across the animal kingdom. Species are not typically defined by molecular distances, so all of the species pairs in our study, though formally described as species, may all be at different genetic stages of divergence, or varied amounts of time of divergence. If all pairs show the same broader pattern of genetic changes, then they can be thought of as being at different stages along the same underlying process. However, if they show very different patterns between species, then multiple underlying processes are possible.

Among congeneric species, many one-to-one orthologous proteins are nearly identical for several pairs (Figure 3A,D, Supplemental Figure 7), and only few proteins were highly divergent (Supplemental Figure 7). Over time, as mutations accumulate throughout the genomes of the two separate species (Figure 3B,C,E,F), the number of completely identical proteins decreases, in a sense turning Figure 3A into 3B. Additionally, the total number of protein pairs being compared also decreases because the more-divergent proteins become difficult to detect by BLAST, effectively “dropping off the graph,” and are neither counted for protein sequence identity nor detected in synteny blocks. The drop-off threshold appears to be at roughly 40% protein identity (Figure 3, see Supplemental Figure 8 for histograms of all species). In the case of *Crassostrea*, where two of the species are now assigned to the genus *Magallana*, the observed distributions of protein identity in Figure 3B,C are potentially a sign of long-past separations, and that what used to be several species in a single genus indeed may be better described as separate genera. Similar distributions of protein identity are seen below in *Drosophila* (Figure 3E,F), and in *Daphnia* (Supplement Figure 8F).

**Figure 3:**
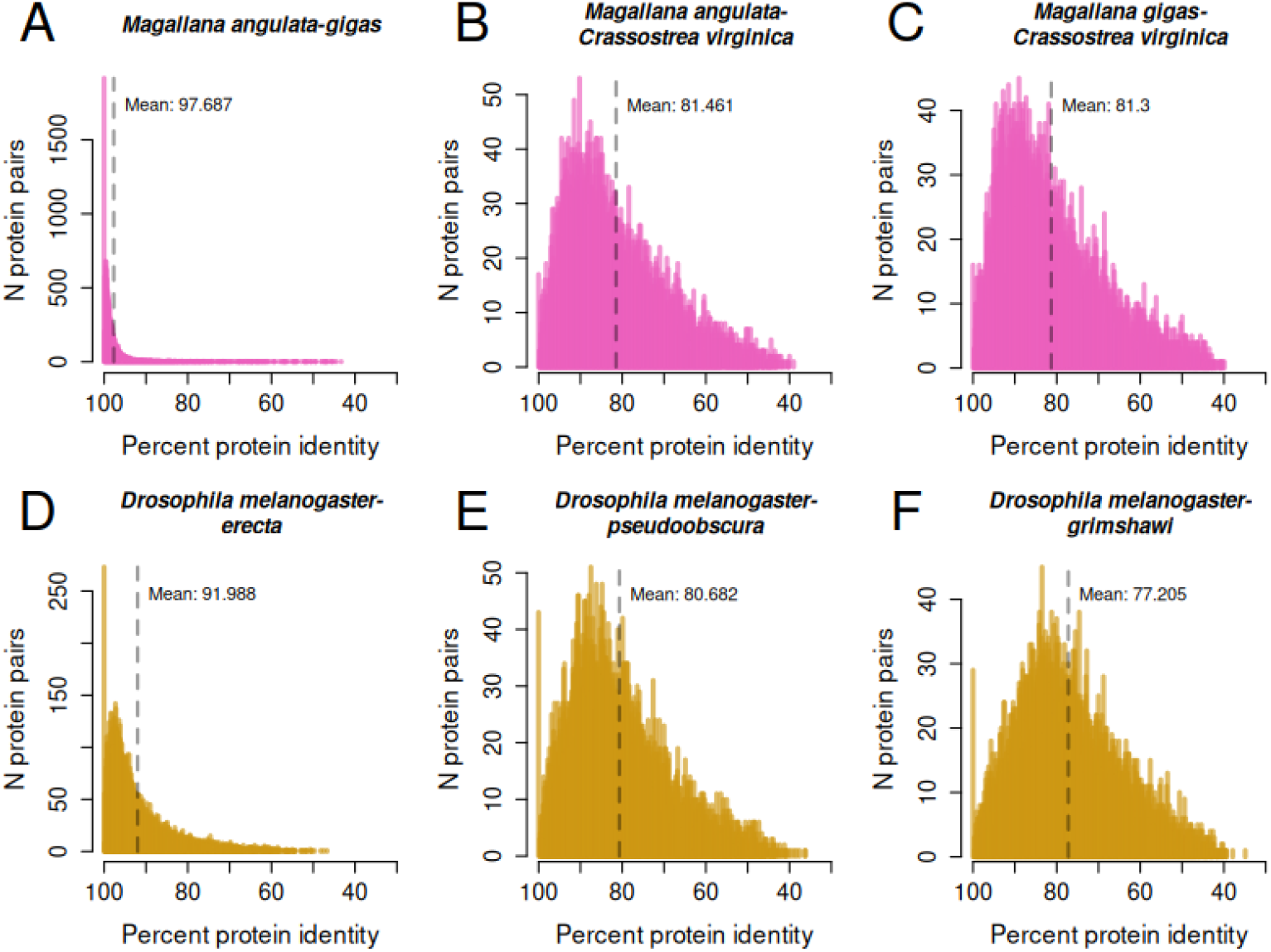
Histograms of protein percent identity for *Crassostrea / Magallana* (A-C) and *Drosophila* (D-F). Protein identity is binned at intervals of 0.1% identity. Note the large differences in Y-axis scale, driven by a large number of 100% identical proteins in A and D.

### Lack of genomic signatures of speciation

The taxonomic hierarchy can be considered an abstraction of the true divergence within a clade. However, taxonomic ranks are arbitrary and do not reflect divergence similarly in different groups. Yet, as ranks can be used as proxies for phylogenetic diversity, taxonomic ranks remain meaningful in contexts such as conservation biology (Hendricks et al., 2014). In our study, as there are multiple trajectories of genetic change among congeneric species (Figure 1), we could not identify a clear genomic distinction based on macro- and/or microsyntenic patterns or percent protein identity that could possibly be useful to delimit different animal species, and generally, any other taxonomic level. Within some clades, patterns related to specific chromosomal features, such as centromere position, may emerge. In our data, some species pairs (e.g., in genera *Crassostrea, Daphnia*) show low levels of microsynteny and relatively low protein sequence identity indicating that the taxonomic rank ‘genus’ in these pairs point to different stages of the evolutionary process (see, for example, also Avise and Liu 2011). For instance, the two oyster species, now *M. angulata* and *C. virginica*, represent a pair where substantial rearrangements are visible in the genome. These two species are also separated geographically, as *M. angulata* was originally from the Pacific, while *C. virginica* is from the Atlantic. The degree of rearrangement in *Daphnia* (Figure 2C) is similar, and potentially this genus should be re-evaluated to see whether it is better described as two genera.

As in most other similar studies, the synteny assignments in our method are ultimately dependent upon local sequence similarity (BLAST, DIAMOND, etc). Hence, one of two effects could result in the exclusion of a gene from our analysis: duplication (no longer a one-to-one ortholog) or mutation beyond the detection of BLAST. This second cause may explain the regression line found by (Zdobnov & Bork, 2007) and the apparent lower bound for protein identity to microsynteny (Figure 1C). In this regard, as protein pairs diverge beyond the detection limit imposed by sequence similarity searches, e.g., in more distantly related species or genes with low evolutionary pressure, they cannot be detected as syntenic. Thus, it is very unlikely to find a species pair with very high microsynteny and low protein identity but not the opposite.

### Difficulty in quantification of syntenic blocks and microsynteny

Meaningful measurements of microsynteny are challenging to get with current data. For instance, assembly quality could affect synteny assessments, as lower quality assemblies would be expected to increase the number of synteny blocks but not necessarily affect the number of syntenic genes. Due to the high sensitivity of the analysis to different parameters, real biological features such as gene insertions, duplications, or long intergenic regions can result in synteny blocks being split in two. Thus, the number of synteny blocks themselves is less important than the number of genes contained in blocks. A further confounding factor is the presence of unassigned scaffolds (i.e., the number of scaffolds is far above the number of chromosomes) in all assemblies analyzed. These may be unmerged haplotypes and would affect the measurement by increasing the number of genes not found in matching synteny blocks as they appear as duplicates.

One attempt to model the syntenic change between mouse and human (Nadeau & Taylor, 1984) had also assumed that reciprocal translocations and not inversions typically caused the synteny breaks. Under that model, it was assumed that the length of synteny blocks follows a negative exponential distribution and that synteny breaks disrupt both micro- and macro-synteny, something not typically observed. Nadeau’s model was disputed by Zdobnov and Bork (2007), who argued that the distribution of block length followed a power law, not an exponential distribution. We found similar correlations in block length in our dataset, though R^2^ ranged between 0.713 and 0.945 (Supplemental Figure 9). Thus, the correlation between block length and number of blocks is not as strong as proposed by Zdobnov and Bork (2007), suggesting that other biological factors affect chromosomal breakpoints (Pevzner & Tesler, 2003).

### Challenges in quantification of rearrangement time

A prominent unanswered question of this study is whether chromosome rearrangements occur at the same rate across a clade. Such a study would be of great interest in evolutionary biology, even if difficult to design correctly. Firstly, such a study would require enough sequenced high-quality chromosome-scale species-pair genomes to identify which lineage had the inversion or translocation based on genomic triangulation. We are currently far from this objective. In our study, there were only 18 genera on NCBI with more than two species with chromosome-level assemblies, and several of those still did not include an annotation.

Secondly, one must infer divergence times between taxa in a way that is not reliant on protein identity. Divergence time estimates are often based on molecular clock inferences. While fossil calibrations are necessary for this analysis, the time estimates are then shifted based on molecular changes, i.e, the inferred divergence time depends on the protein identity. Unsurprisingly, one would likely find a strong correlation between protein identity and divergence time (as in Supplemental Figure 10), but that is a problem of circular logic; the divergence time was a function of protein identity in the first place, so it cannot be applied as an independent measurement for rearrangements. This is extended with the correlation of protein identity to the number of genes in synteny blocks. Consequently, protein identity, microsynteny, and molecular clock-based divergence estimates are likely correlated. Unfortunately, this means that molecular clock divergence time should not be used to test the rearrangement rate in lineages. One possibility for timing narrow-scale speciation could be the usage of codon evolution models. For instance, in Placozoa, it was observed that the nucleotide identity differed from protein identity such that highly conserved genes had many synonymous nucleotide changes instead of simply having fewer total mutations (Eitel et al., 2018). Thus, codon evolution models could provide a different age estimate than the proteins.

## Conclusions

Defining and delimiting species is one of the important and basic requirements for studying biodiversity and its evolution, and it is pivotal for the management and conservation of natural resources. However, even in the times of DNA barcoding and genome sequencing, no universal threshold that applies, for example, across all animals, has been identified, and different species concepts are applied in different taxa. Synteny, i.e., the shared arrangement of genes on chromosomes between species, has been used as a marker of shared ancestry, with synteny-breaking events resulting in genomic incompatibilities between populations, ultimately leading to speciation. However, in our study of the genomes of congeneric pairs, we found that genomic changes inferred from speciation are variable and contextual and that synteny should not be used at present as a measure of divergence or speciation. Once more high-quality, chromosome-scale genomes of sister species-pairs are available, a clear pattern of genome evolution may emerge.

## Data, script, and code availability

Additional data, figures, scripts and analytical resources can be accessed in a GitHub repository https://github.com/PalMuc/congeneric_synteny, archived with a DOI at Zenodo (https://zenodo.org/doi/10.5281/zenodo.10069099).

## Supplementary Information

Supplementary information is available from https://github.com/PalMuc/congeneric_synteny/tree/main/supplements_for_paper

## Acknowledgements

WRF would like to thank M. Dohrmann and D. Schultz for helpful discussions. We would like to thank M. Eitel for helpful discussions and comments. G.W. dedicates this work to his late wife Connie Wörheide, who left us too early during the preparation of this manuscript.

