## Supplemental Figure for "Genomic changes are varied across congeneric species pairs of animals"

### Supplemental Figures

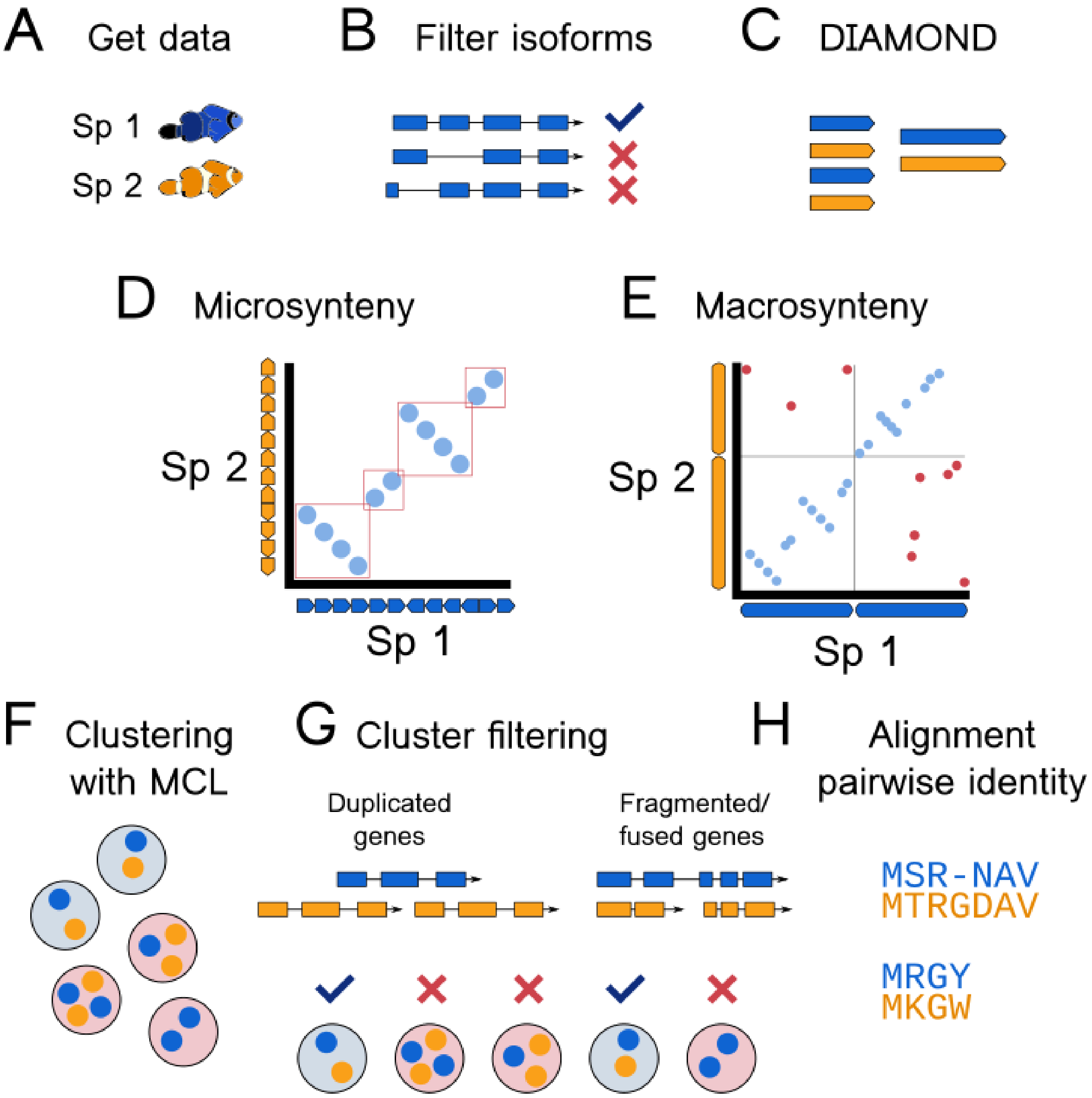


Supplemental Figure 1: Schematic of analysis (A) Chromosome-level genome assemblies and associated data are used for two congeneric species. (B) Data were filtered to remove splice variants. (C) Align all proteins from both species using DIAMOND, used for steps D, E and F. (D) Identify microsyntenic blocks (E) Identify overall macrosynteny. (F) Cluster proteins with mcl. (G) Filter clusters to include only one-to-one orthologs, making sequence pairs. (H) Align all sequence pairs using MAFFT, used for further calculations.

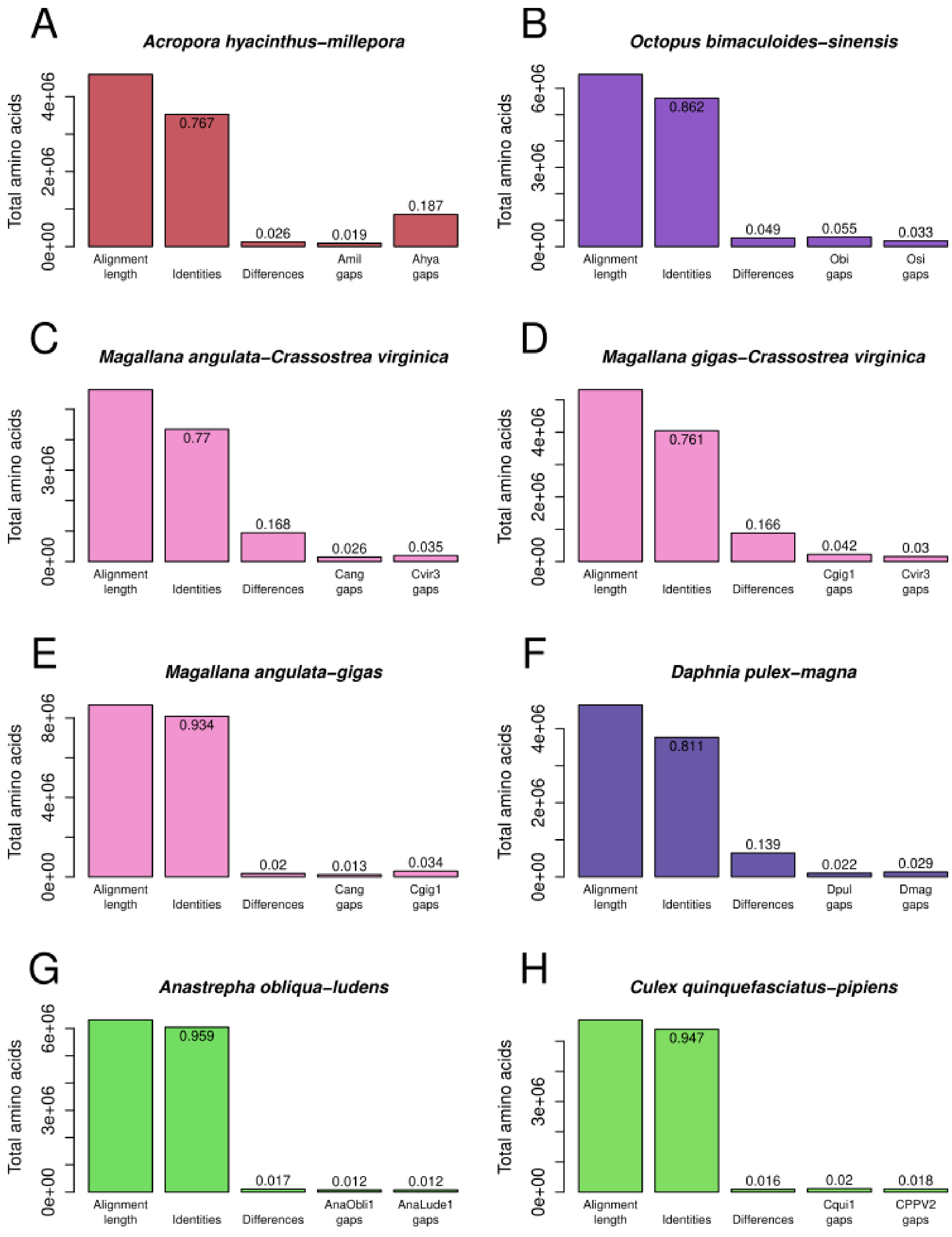


Supplemental Figure 2: Comparison of all pairwise protein alignments across all species pairs (A-H).

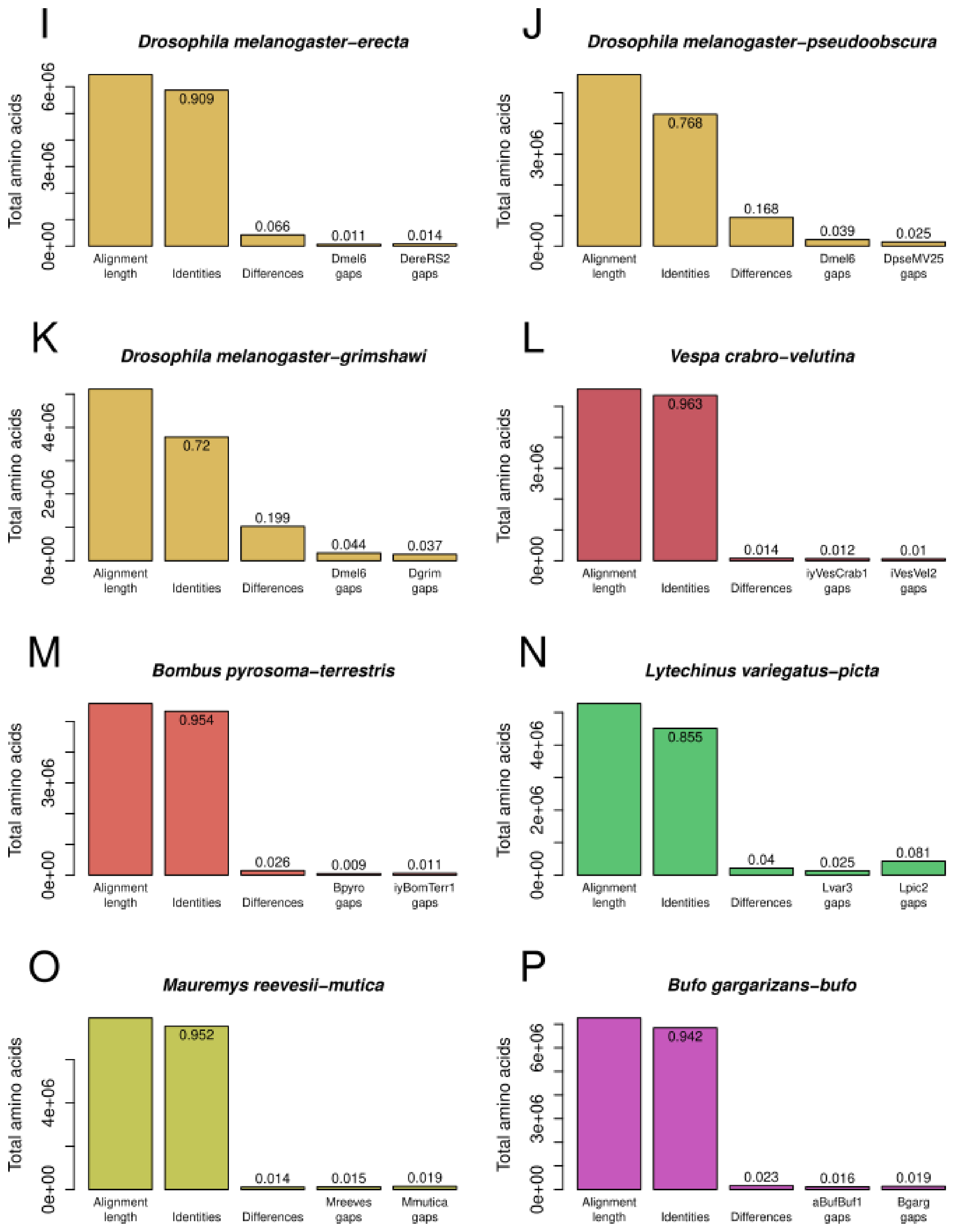


Supplemental Figure 2 continued: Comparison of all pairwise protein alignments across all species pairs (I-P).

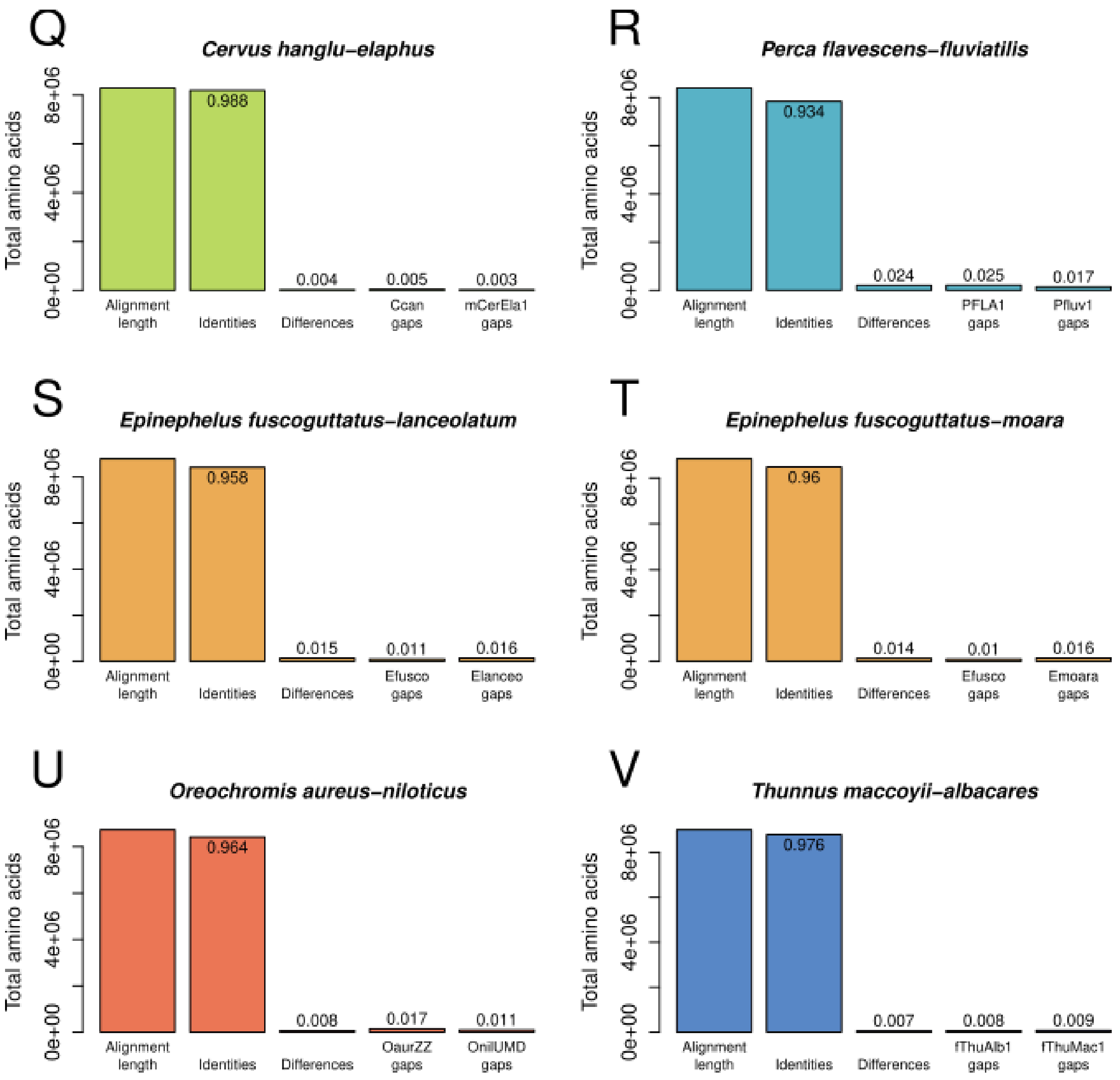


Supplemental Figure 2 continued: Comparison of all pairwise protein alignments across all species pairs (Q-V).

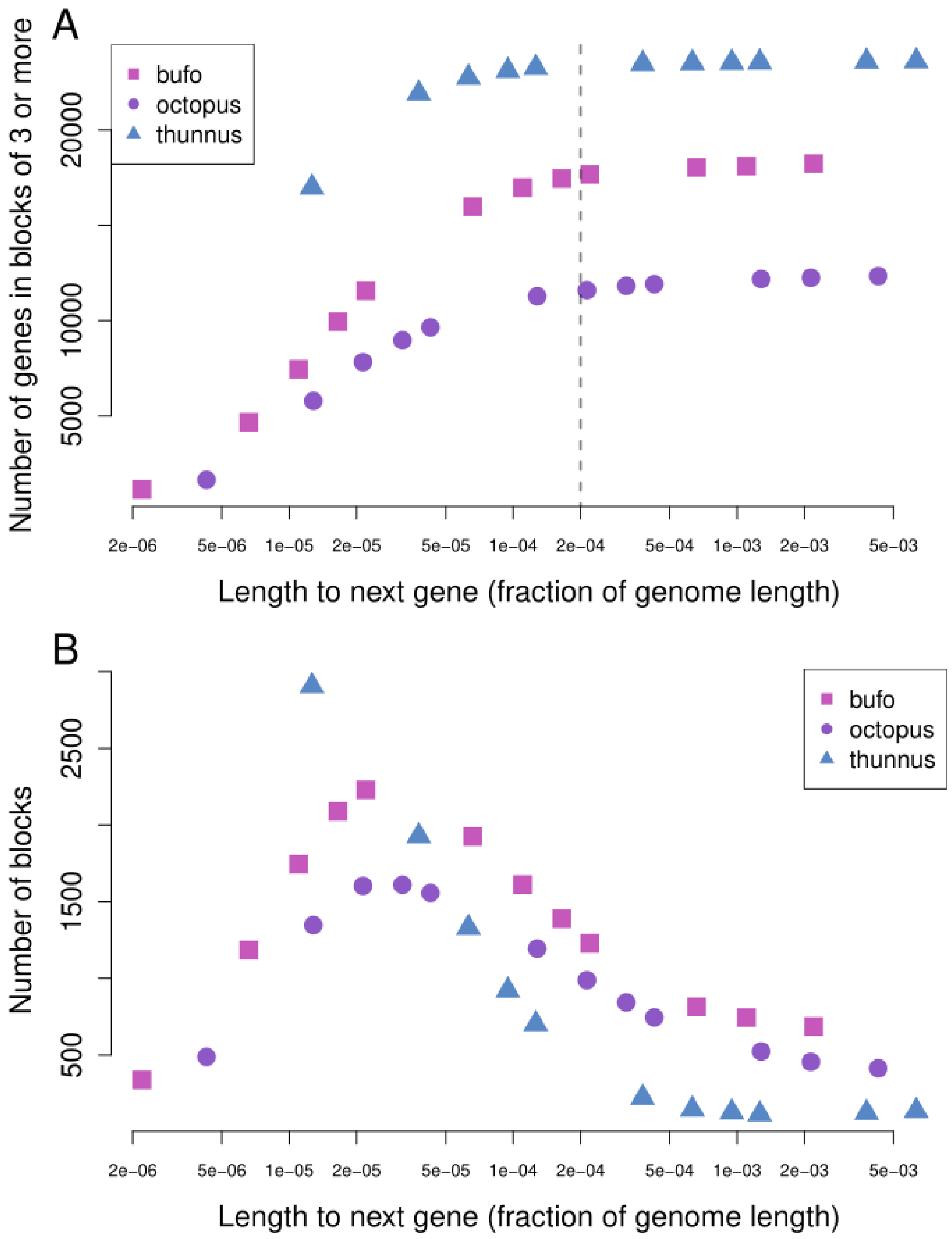


Supplemental Figure 3: Sensitivity of the detected gene blocks to the skip parameter **-z** for three genera with large genomes, *Bufo* (appx. 4.5Gb), *Octopus* (appx. 2.5Gb) and *Thunnus* (appx. 800Mb). Skip parameter values were given as length in bp, but were normalized by the average genome length for each genome pair. (A) Effect on the number of recovered microsyntenic genes. The gray line indicates $2x10^{-4}$ (1/5000) times the total genome size, which shows little increase in microsyntenic genes beyond this value. (B) Effect on total number of blocks. With increased **-z**, smaller blocks become joined into large blocks, reducing the total number of blocks.

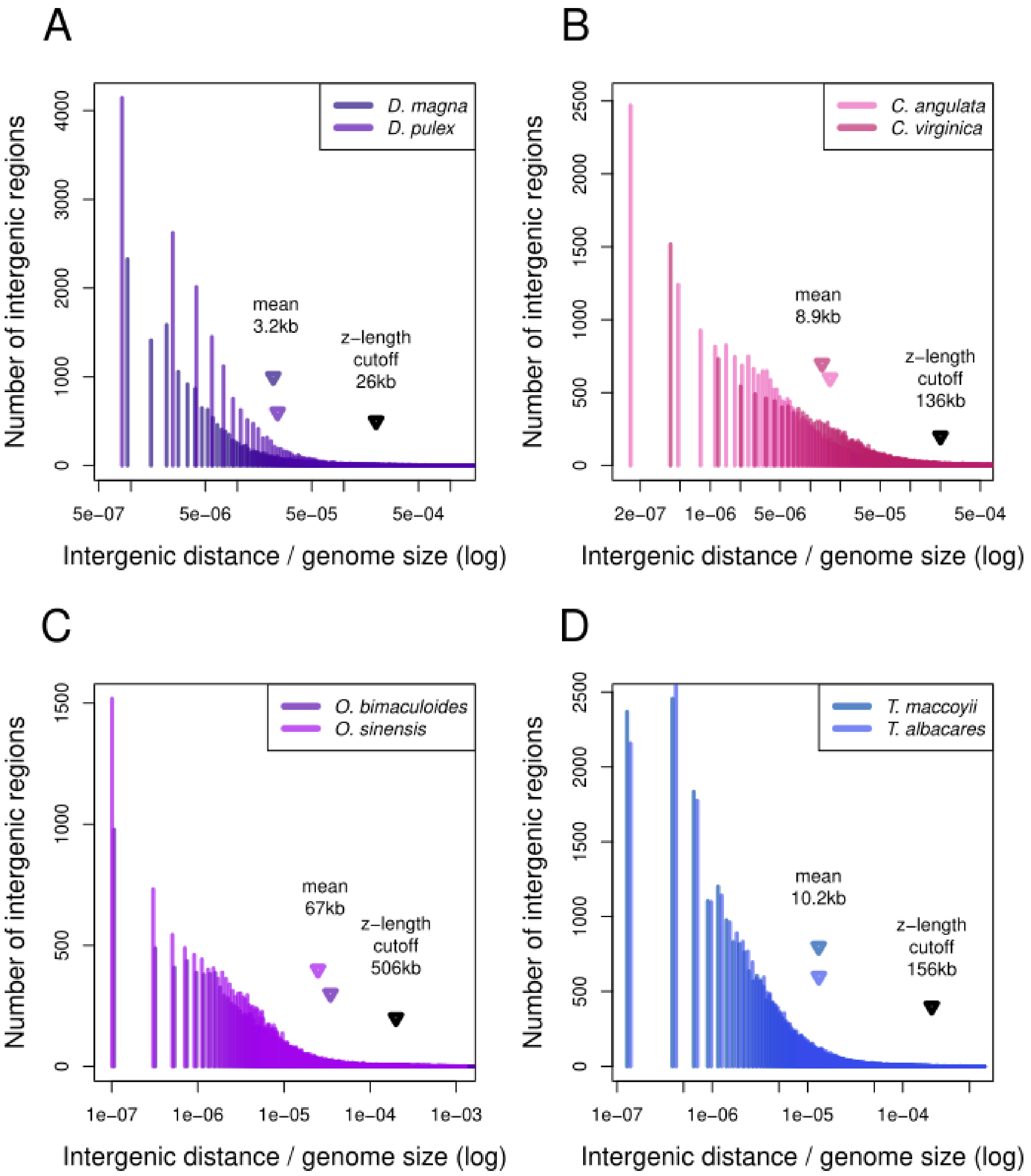


Supplemental Figure 4: Histogram of intergenic distances for *Daphnia*, *Crassostrea*, *Octopus*, and *Thunnus* (A, B, C and D, respectively). Intergenic lengths are normalized by the genome size, to correspond to Supplemental Figure 2. The vast majority of intergenic distances are shorter than the calculated skip parameter **-z** for each species pair (corresponding to 1/5000 times the genome size).

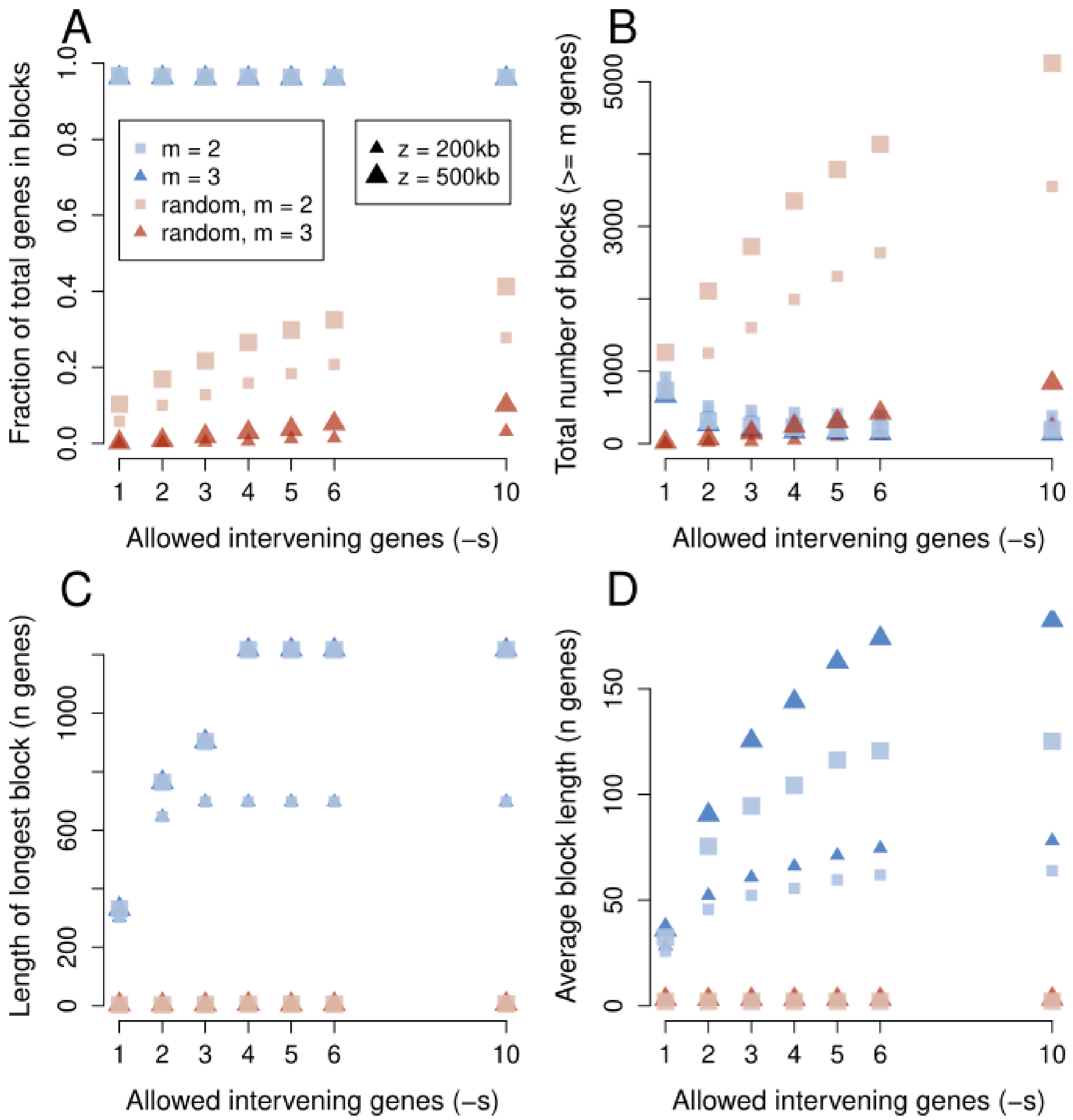


Supplemental Figure 5: Combined effects of number of allowed intervening genes, for real (blue) and randomized (red) data from *Thunnus*. For all panels, parameter **-m** indicates the minimum allowed block size, either 2 or 3 genes, and parameter **-s** indicates the number of allowed intervening genes. (A) Fraction of total genes recovered in syntenic blocks. (B) Total number of microsyntenic blocks. (C) Length of longest block, in genes. (D) Average block length, in genes. Panels A and B show that most spurious matches occur in randomized data when the minimum block length is set to 2, and increase with the number of intervening genes. This does not strongly affect real data.

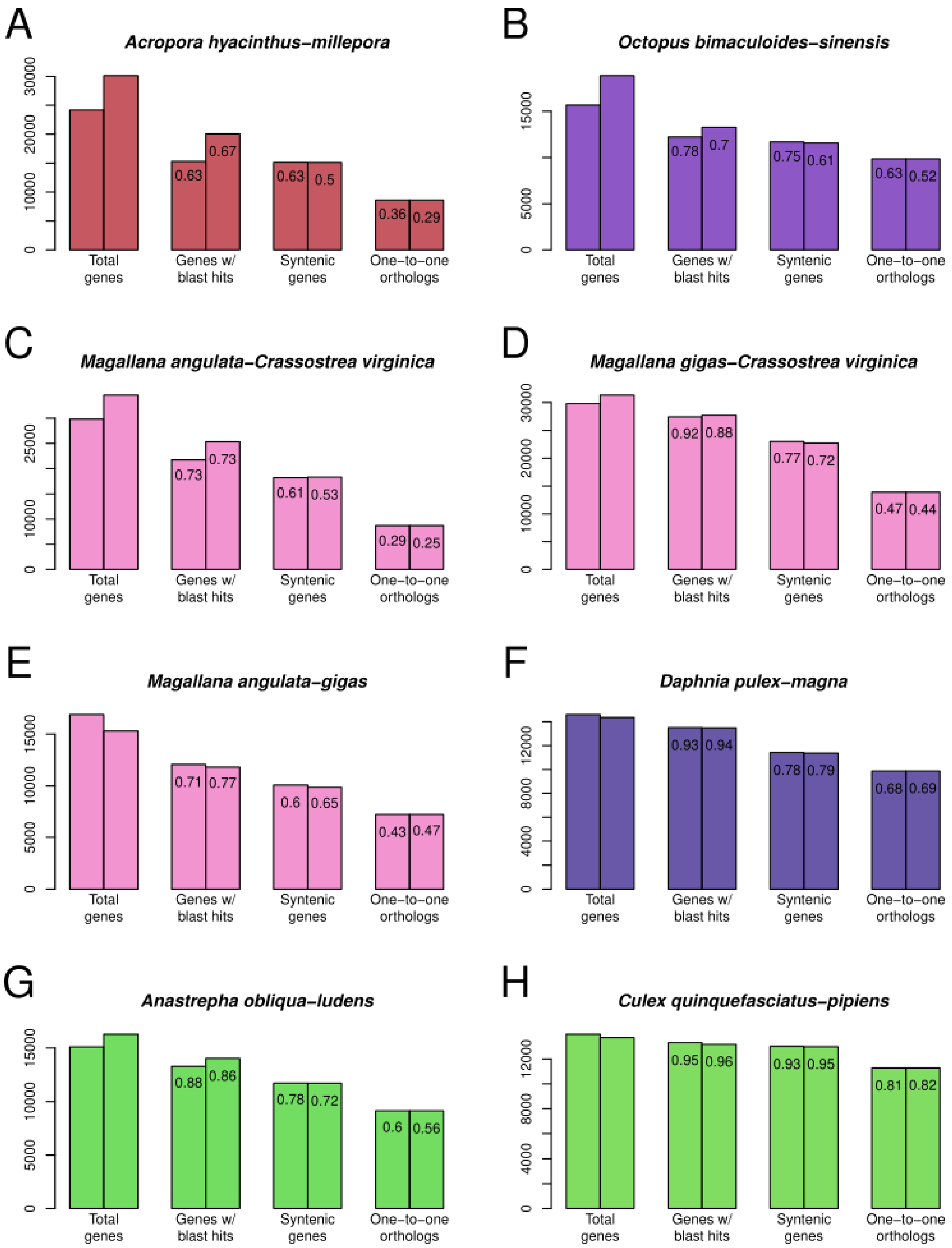


Supplemental Figure 6: Counts of protein pairs used for protein alignments for all species pairs (A-H).

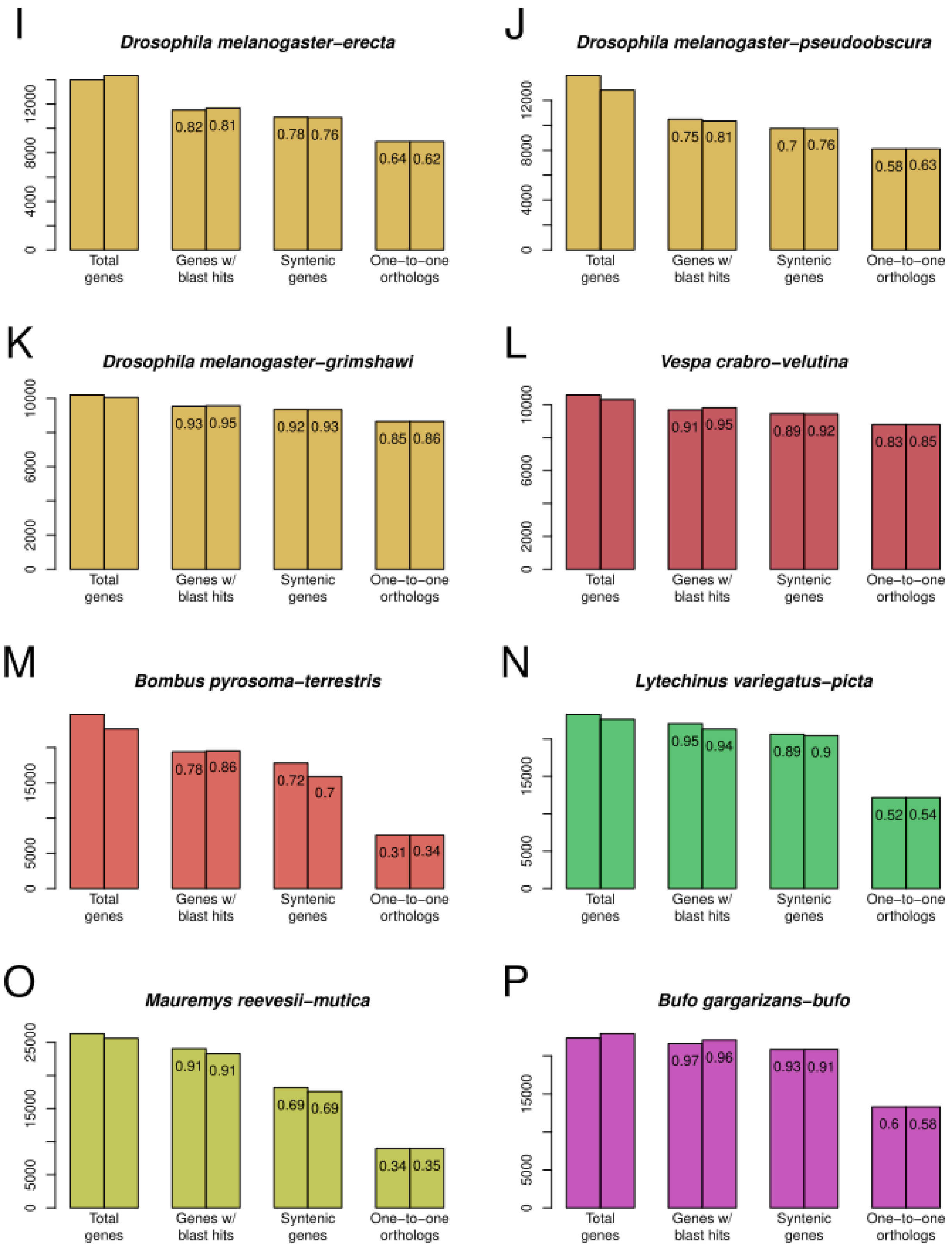


Supplemental Figure 6 continued: Counts of protein pairs used for protein alignments for all species pairs (I-P).

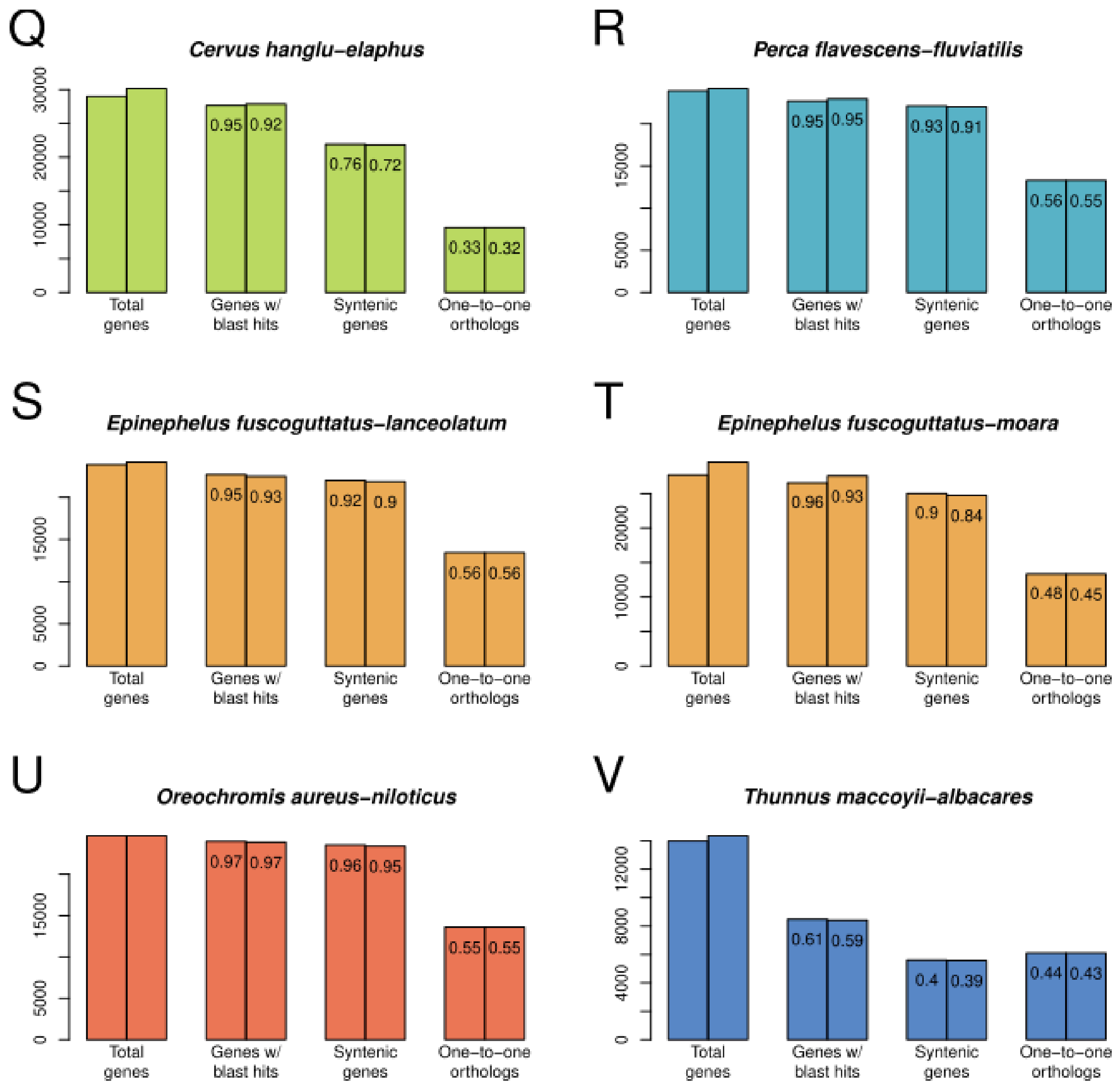


Supplemental Figure 6 continued: Counts of protein pairs used for protein alignments for all species pairs (Q-V).

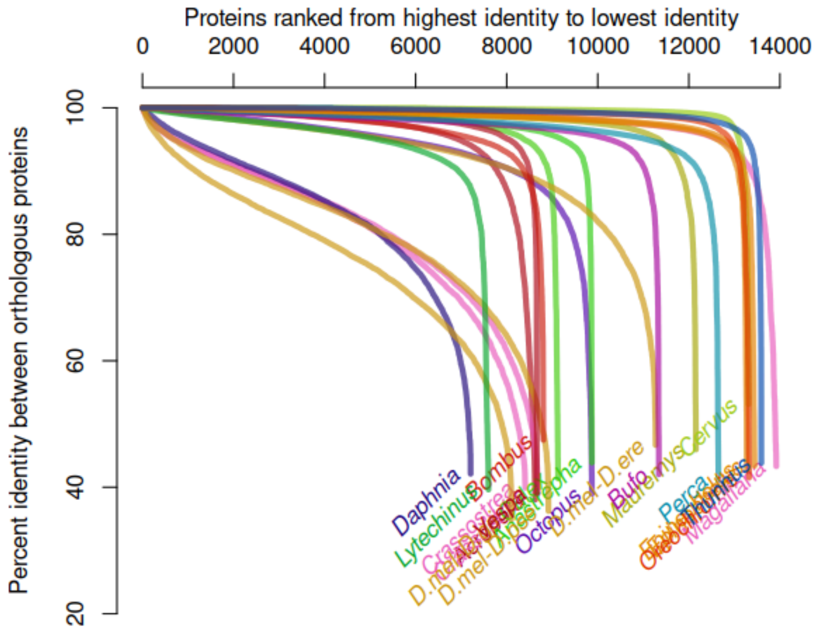


Supplemental Figure 7: Protein identity of orthologous proteins ranked from highest to lowest for each species. The majority of proteins are nearly identical for many species (forming the horizontal lines near the top of the graph) followed by few highly divergent proteins (the vertical portions of the lines). Many labels are moved for clarity.

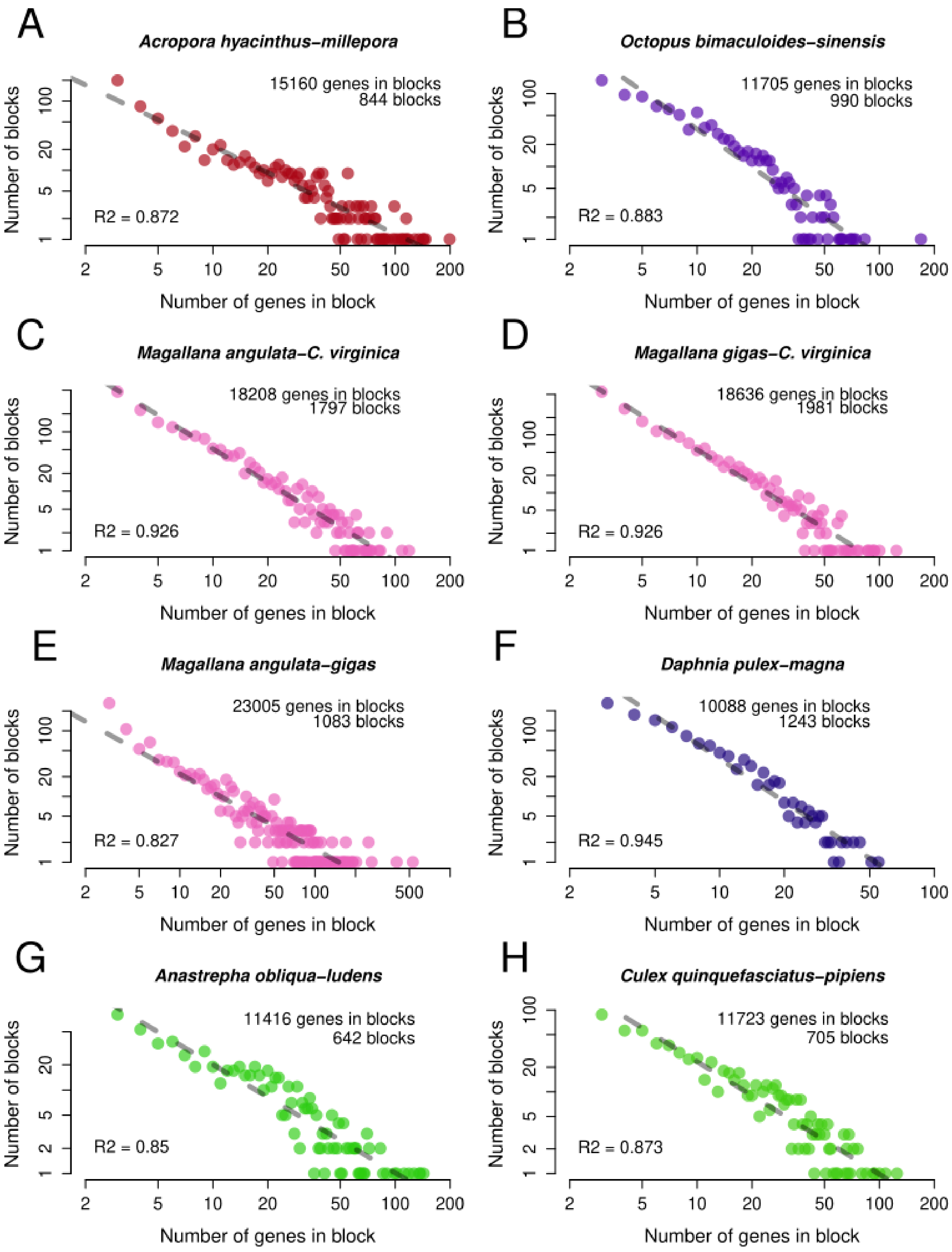


Supplemental Figure 8: Histogram of pairwise protein sequence identities, across all species pairs (A-H).

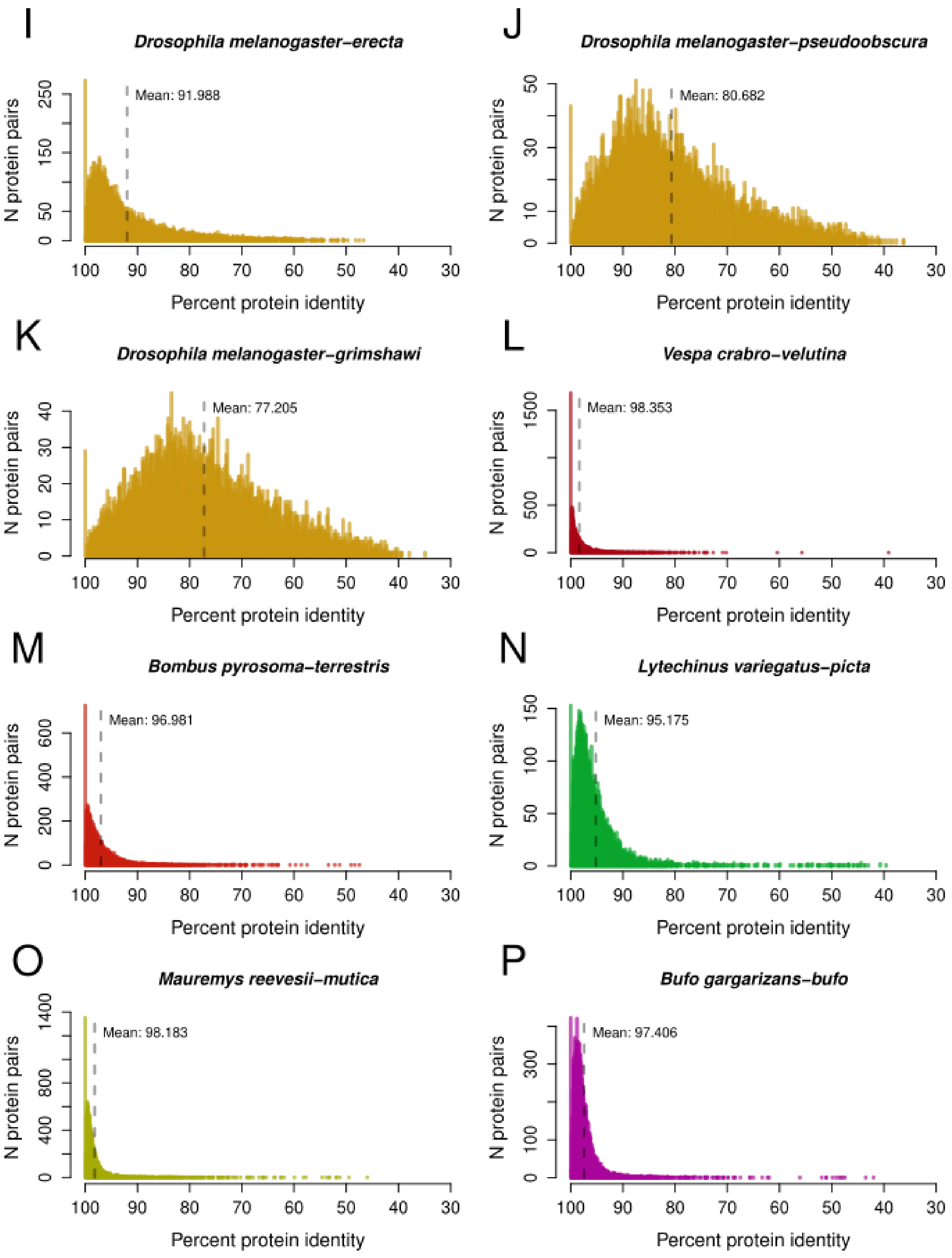


Supplemental Figure 8 continued: Histogram of pairwise protein sequence identities, across all species pairs (I-P).

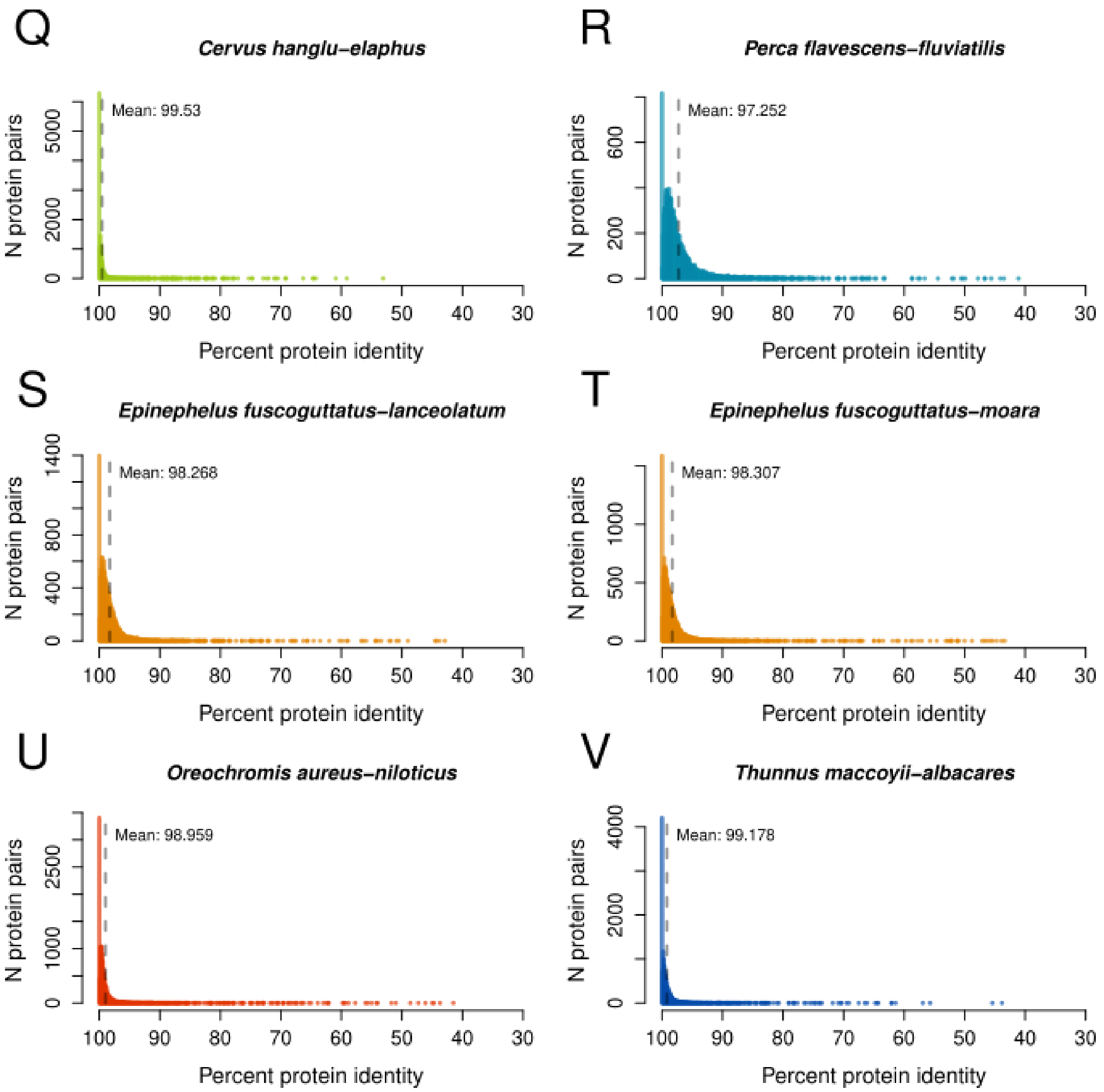


Supplemental Figure 8 continued: Histogram of pairwise protein sequence identities, across all species pairs (Q-V).

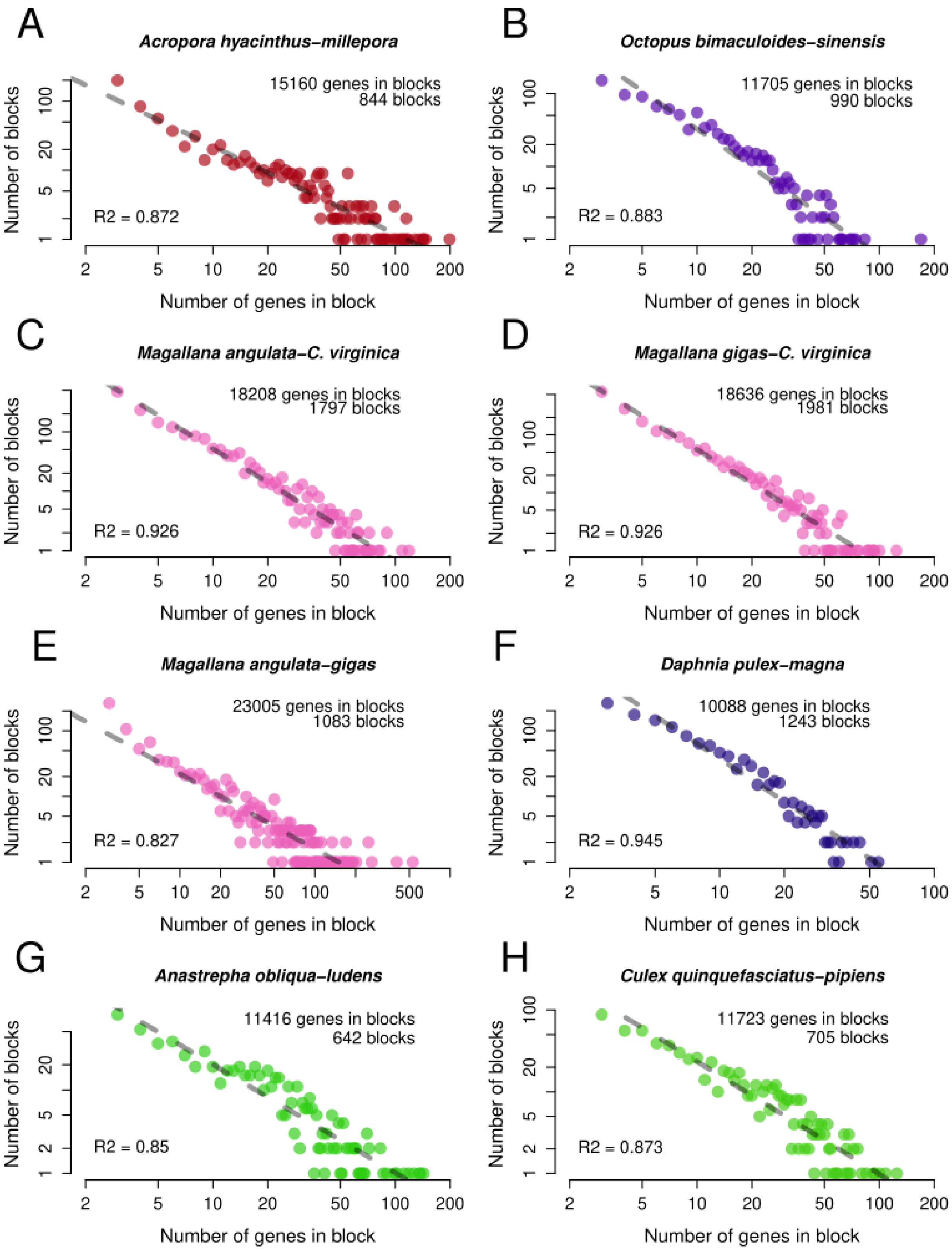


Supplemental Figure 9: Counts and lengths of microsynteny blocks, across all species pairs (A-H). The proposed power law relationship (log-log) is visible in some pairs. The linear regression is indicated as the dotted gray line for all species pairs.

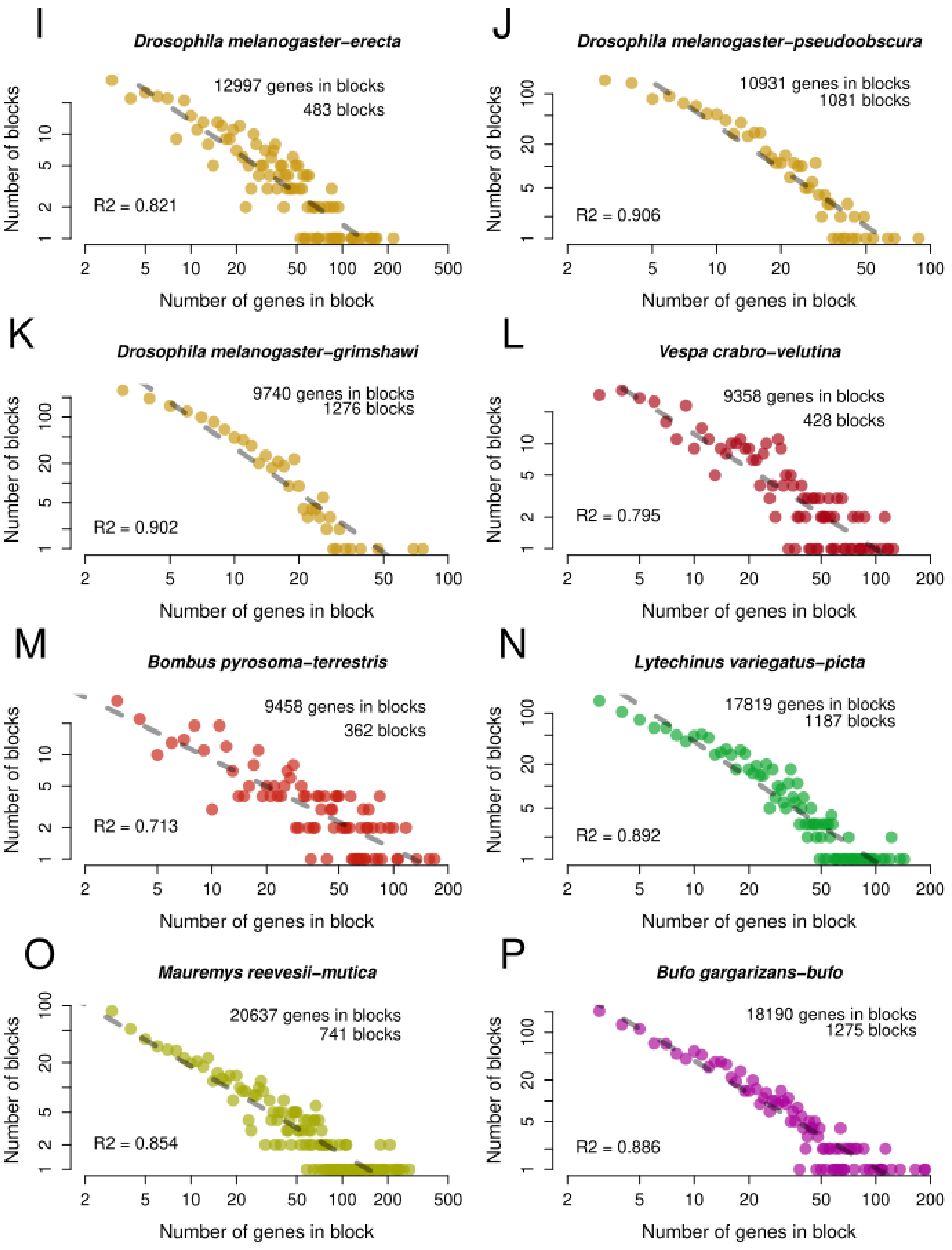


Supplemental Figure 9 continued: Counts and lengths of microsynteny blocks, across all species pairs (I-P).

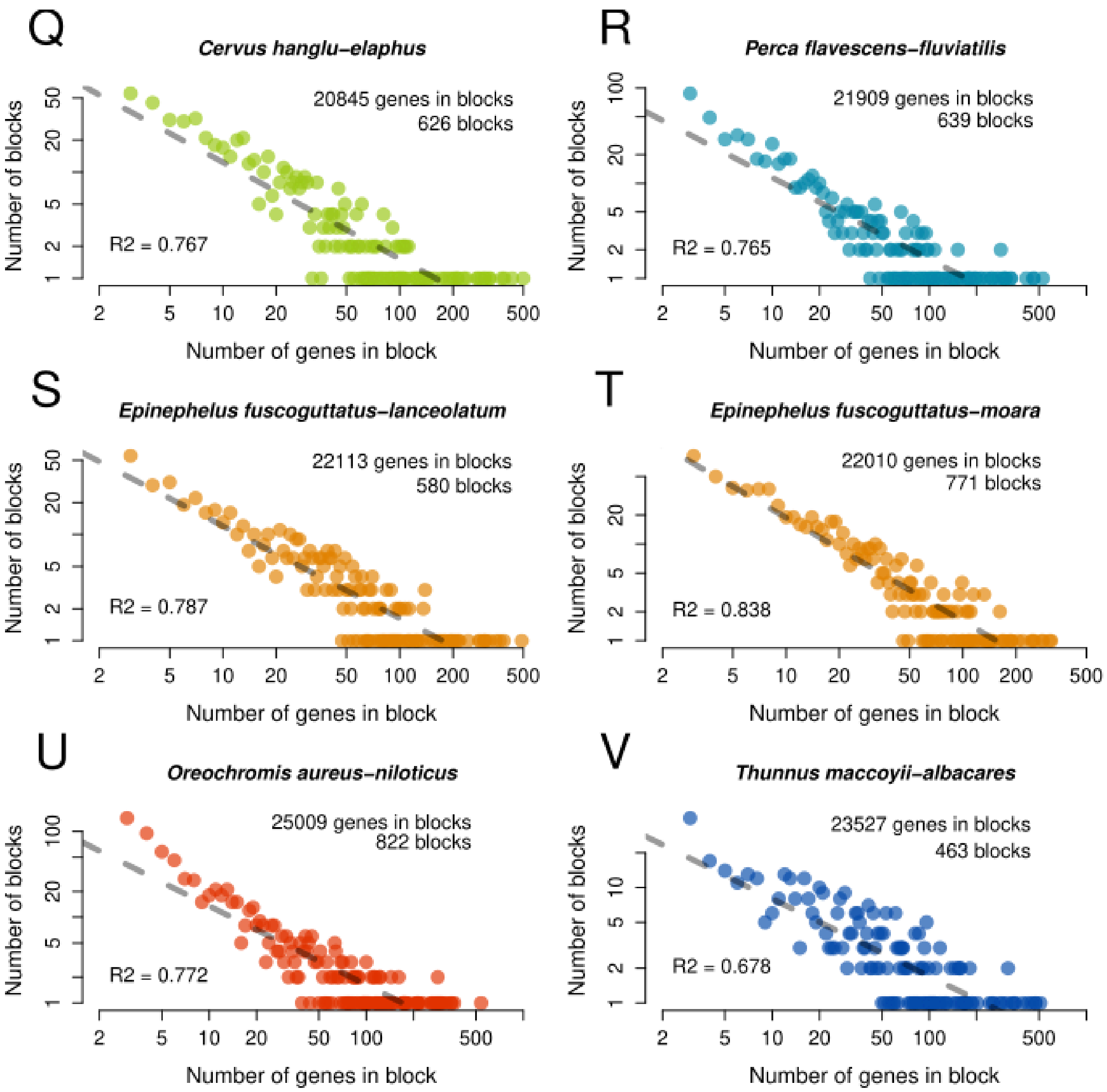


Supplemental Figure 9 continued: Counts and lengths of microsynteny blocks, across all species pairs (Q-V).

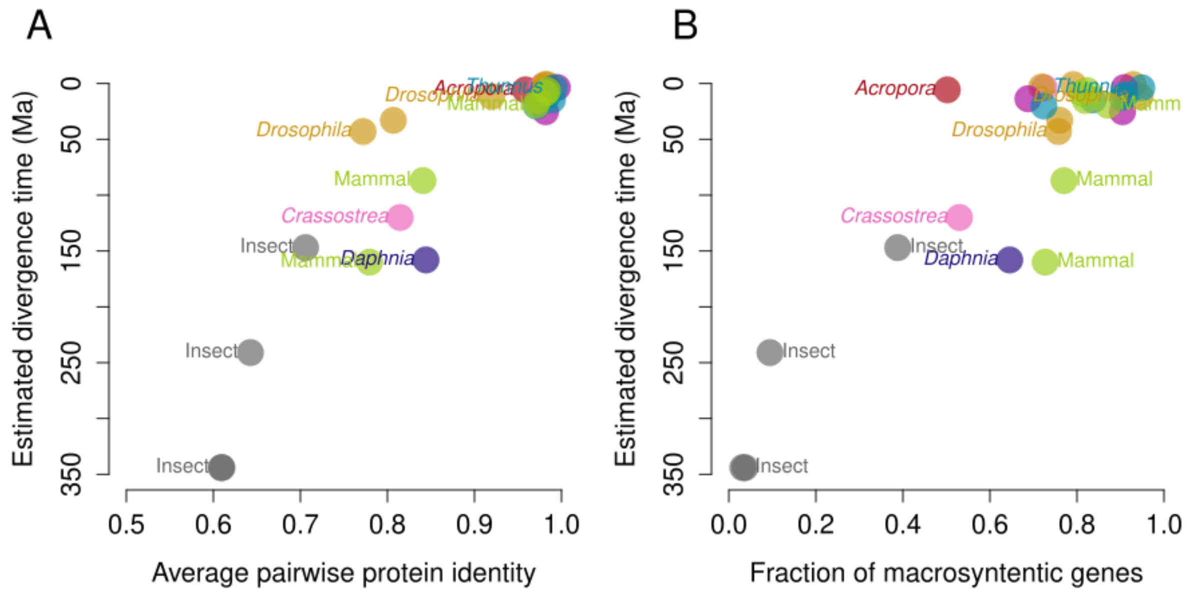


Supplemental Figure 10: Protein identity (A) and macrosyntenic genes (B) compared against divergence times taken from timetree.org, where available.
